# Purging genomes of contamination eliminates systematic bias from evolutionary analyses of ancestral genomes

**DOI:** 10.1101/2022.11.17.516887

**Authors:** Balázs Bálint, Zsolt Merényi, Botond Hegedüs, Igor V. Grigoriev, Zhihao Hou, Csenge Földi, László G. Nagy

## Abstract

Contamination of genomes and sequence databases is an increasingly recognized problem, however, efficient tools for removing alien sequences are still sparse and the impact of impure data on downstream analyses remains to be fully explored. Here, we present a new, highly sensitive tool, ContScout, for removing contamination from genomes, evaluate the level of contamination in 844 published eukaryotic genomes and show that contaminating proteins can severely impact analyses of genome evolution. Via benchmarking against synthetic data, we demonstrate that ContScout achieves high specificity and sensitivity when separating sequences of different high level taxa from each other. Furthermore, by testing on manually curated data we show that ContScout by far outperforms pre-existing tools. In the context of ancestral genome reconstruction, an increasingly common approach in evolutionary genomics, we show that contamination leads to spurious early origins for gene families and inflates gene loss rates several fold, leading to false notions of complex ancestral genomes. Using early eukaryotic ancestors (including LECA) as a test case, we assess the magnitude of bias and identify mechanistic bases of the estimation problems. Based on these results, we advocate the incorporation of contamination filtering as a routine step of reporting new draft genomes and caution against the outright interpretation of complex ancestral genomes and subsequent gene loss without accounting for contamination.

## Introduction

Recent technological advances in high-throughput sequencing and plummeting sequencing costs are leading to unprecedented growth in genomic sequence databases^1,2^. Instruments that deliver long-read sequences and the vastly improved throughput of short-read platforms are enabling the resolution of complex eukaryotic genomes in addition to the prokaryotic ones that dominated early sequencing projects. Recently, several large-scale eukaryote sequencing projects have been launched with the goal of capturing the genomes of tens of thousands of insects^3^, vertebrates^4^, fungi^5^, plants^6^, or ultimately the entire eukaryotic biodiversity on Earth^7^. Notably, unlike many completely resolved (finished) prokaryote genomes, nearly all eukaryotic genomes enter public databases as unfinished drafts.

Due to various biological or technical issues, draft genomes may contain sequences that are not part of the targeted organism^8,9^. Symbionts or pathogens associated with the target could serve as biological sources of contamination, whereas technical sources include sample mishandling and data processing errors. Projects that rely on museum specimens are particularly vulnerable to cross-contamination^10–12^. If not carefully addressed, contaminated reference genomes can poison public databases with inaccurately labeled sequence data, as demonstrated by a recent study that identified over 2 million contaminated records in GenBank alone^13^. The extent of contamination within a draft genome can vary from project to project, but in some extreme cases a near-complete draft of the contaminant organism’s genome can be assembled from the sequencing data in addition to the one of the targeted specimen^14,15^.

It is known that contamination can compromise downstream analyses, be misinterpreted as horizontal gene transfer ^16,17^ and can negatively affect phylogenetic tree inference ^18–20^. However, the sensitivity of other phylo- and evolutionary genomic approaches to contamination has not been assessed, despite significant expansion of these fields^21,22^. For example, ancestral genome estimation was used to reconstruct genomes of extinct organisms *in silico^23–25^*, infer the genome organization and complexity of ancestral organisms^26–28^, gene duplication and loss patterns^21,22^, rates of genome evolution^29^ and the genetic underpinnings of major evolutionary transitions^30,31^, among others. This framework has also been used to predict genes involved in a trait of interest by correlating phenotypic with genetic changes (forward genomics)^32,33^. Although a sensitivity of these approaches to certain analytical issues has been documented (e.g. Pett et al.^34^, Hahn^35^) whether and how data quality issues, such as contamination affect their performance has not been examined.

In the last decade, several tools were developed to detect contamination, based on a range of search logics, such as database-free or -dependent taxonomic classification of raw reads or genes using BLAST searches or k-mers, or utilizing either pre-selected marker genes or genome-wide catalogs to assessing genome quality. However, all of these approaches have limitations that preclude their use for explicit tagging and removal of contaminating genes/proteins. Tools that build on selected universal single-copy genes (e.g. CheckM^36^, BUSCO^37^ and ConFindr^38^) can accurately estimate the extent of contamination, but can not identify and remove the exact contaminating sequences. Most contamination assessment tools, such as CheckM^36^, CLARK^39^, ConFindr^38^ Anvi’o^40^ and GUNC^41^ focus exclusively on prokaryotes (Archaea, Bacteria) or accept only DNA sequences as input (e.g. Kraken^42^, ProDeGe^43^, BlobTools^44^, PhylOligo^45^, CroCo^46^, CONSULT^47^. Because DNA evolves faster than protein sequence, tools that use the former implicitly assume that the contaminating organism, or its close relative is present in reference databases. This is often not the case even in the best-sampled organismal groups, suggesting that protein based solutions may be necessary, especially in the context of comparative phylogenomic studies which usually use protein sequences as input. BASTA^48^ and Conterminator^13^ can both use protein sequences as input, and the latter was used to flag over 2.1 million genes in RefSeq and ~14,000 proteins as contamination in NR. Despite these developments, efficient and highly sensitive tools that can flag and remove contaminating proteins from genome databases are currently lacking.

In this paper we present a new, accurate tool, ContScout, for identifying and removing contaminating proteins from draft proteomes. Rather than applying a fixed threshold, ContScout automatically selects the top-scoring hits for each query sequence that are optimal for taxonomic classification and combines this information with gene position data, yielding improved classification accuracy over pre-existing methods. Screening 844 published eukaryotic genomes with ContScout, we identify 51,222 contaminating sequences originating mostly from bacteria but also from fungi, metazoans and plants. We further show that the inclusion of contaminating sequences in evolutionary genomic analyses leads to spurious ancestral gene count estimates and severely inflated gene loss rates. We demonstrate the adverse effects of contamination on ancestral genome composition of key eukaryotic ancestors including LECA.

## Results

### Novel algorithm for the detection and removal of contamination from draft genomes

Here, we developed a novel contamination detection and removal tool, ContScout that combines reference database-based protein taxonomy classification with gene position data (Fig. 1). Each predicted protein from a query genome is first associated with a high-level taxon (HLT) tag via a speed-optimized protein sequence search against a taxonomy-aware reference database using either DIAMOND^49^ or MMseqs^50^.

**Figure 1:**
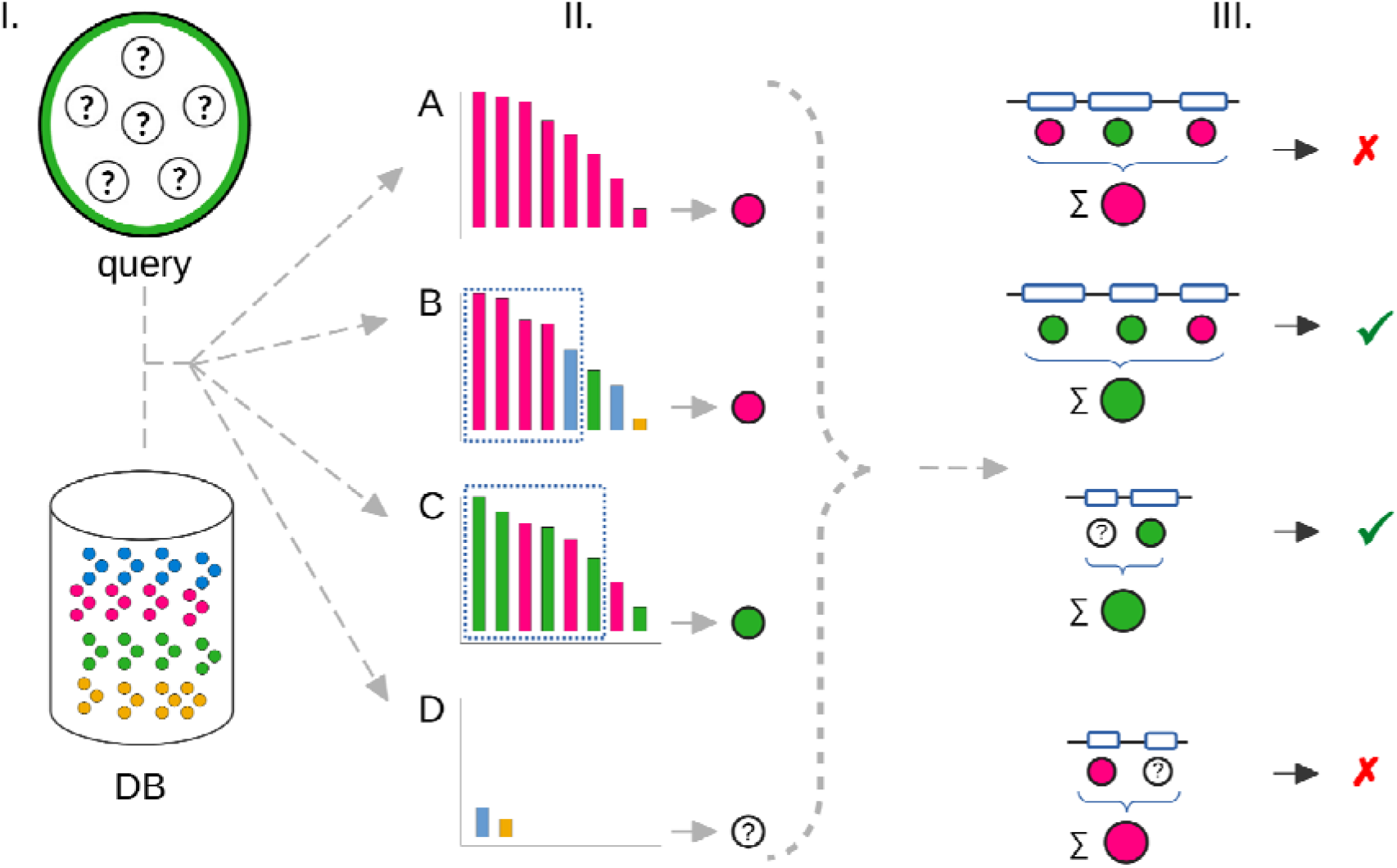
Overview of the ContScout algorithm. **I**. A rapid database search with the query proteins is carried out against a taxonomy-aware reference database. **II**. Hit trimming with a dynamic threshold is carried out for each hit list and a high level taxon label is assigned for each of the query proteins. **III**. Protein taxon labels are summed over coding contigs / scaffolds. Whenever a contig / scaffold disagrees with the query genome, it is removed, together with all the proteins it encodes. Protein taxon call examples: **A**., all hits agree: no dynRLE trimming is applied. **B**., Hit list contains more than two high level taxa: top-scoring hits up to two taxa are considered. **C**., Hits from two taxa alternate: up to three taxon changes are allowed. **D**., Insufficient number of hits observed: protein is labeled as “unknown”. For examples **B** and **C**, blue dotted squares indicate top hits that are taken into account upon query protein taxon labeling. Green check marks indicate genuine host sequences while red cross marks indicate sequences marked as contaminants.

Hit lists with reference HLT labels are then trimmed retaining only the top-scoring upper part with up to two HLT tags and with no more than three value changes in total (Fig.1, part II.). In order to increase sensitivity and specificity, HLT classifications are then combined with coding site information (contig / scaffold annotations) leading to consensus HLT labels over each contig / scaffold in the assembly. Contigs with HLT labels matching that of the query proteome are kept while those ones that disagree are marked as contamination and are removed with all proteins they encode.

The data storage footprint of ContScout is between 0.1-7,8 GBytes per query genome with a run time of 46-113 minutes benchmarked using 24 CPU cores with the RAM usage being constrained to 150 GB. The rate-limiting step is the similarity search step that accounts for 80-99% of the total run time. (Supplementary Fig. 1).

ContScout is implemented in R, with all software components and their dependencies placed in a Docker container for easy deployment. The software package also contains a database downloader script that allows for a convenient download and pre-formatting of reference databases.

### Performance assessment on synthetic data

Performance of ContScout was assessed by creating artificially contaminated genomes for all possible pairwise cross-HLT pairs of 17 contamination-free genomes from the G36 data set (see methods for details). For each genome pair, both contamination directions were tested by introducing 100, 200, 400, 800, 1600 or 3200 contaminant proteins in the receiver proteome.

Testing with synthetic data revealed that ContScout overall identified the vast majority of contaminating proteins in each combination (Fig. 2). It performed best when the source of contamination came from Bacteria (AUC range: 0.98-1, median: 1), Animals (AUC range: 0.9649-1, median: 0.99), Plants (AUC range: 0.98-1, median: 0.99) or Fungi (AUC range: 0.97-1, median: 0.99). Somewhat decreased performance was observed when other eukaryotes, representatives of Euglenozoa and Amoebozoa were used as the source of contamination (AUC range: 0.64-1, median 0.98). In line with the lower AUC values, slightly decreased true positive rates (TPR range: 0.78-1, median: 0.98) were observed when *Phytomonas* was the source of contamination while contaminating with *Dictyostelium* sequences resulted in the worst true positive rate (TPR range: 0.28-1, median: 0.82) regardless of the target genome tested.

**Figure 2:**
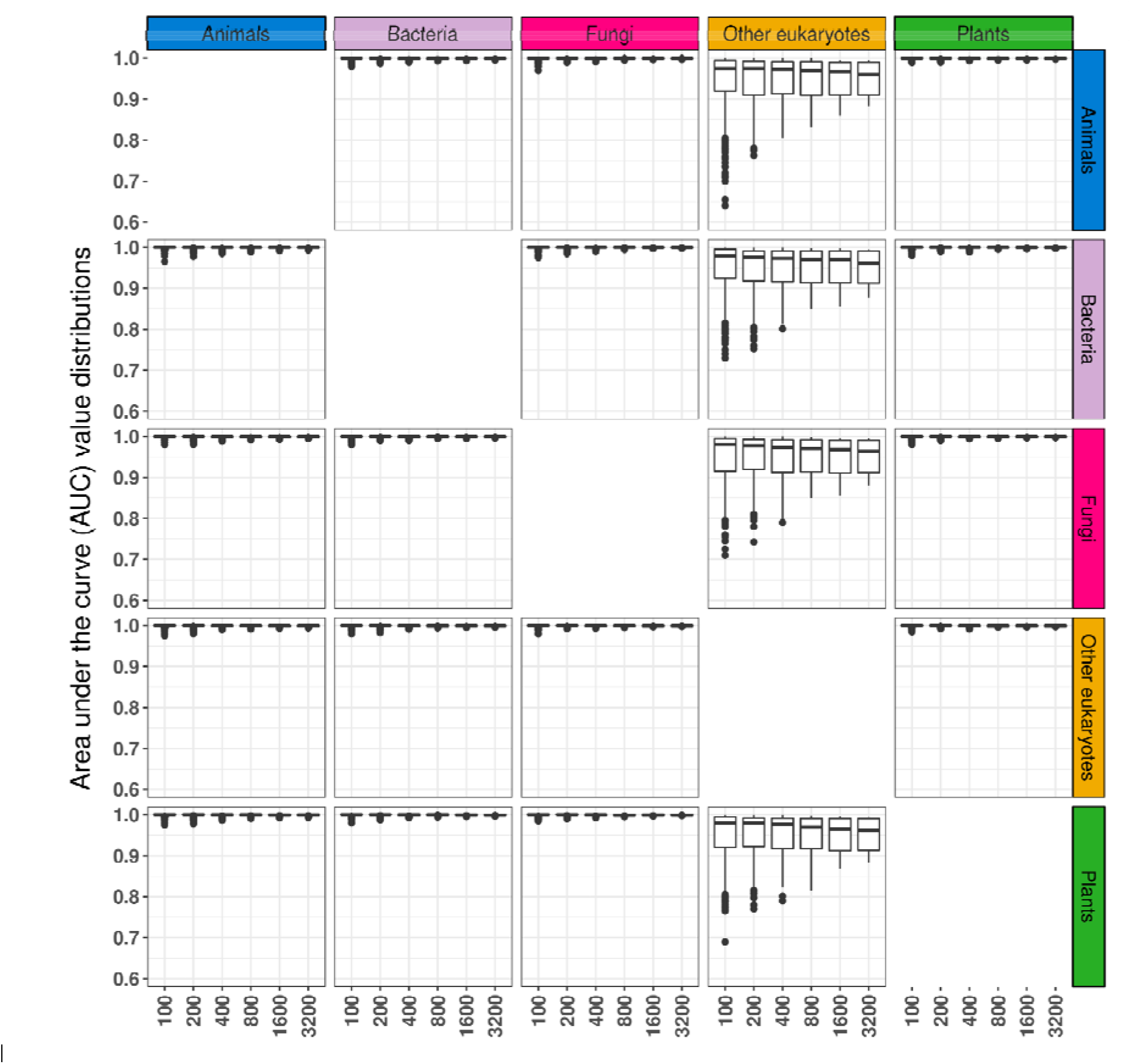
Performance of ContScout on synthetic data. Artificially contaminated genomes were created by transferring varying numbers of proteins between all possible combinations of source and target proteomes from various HLTs. Artificially introduced proteins were classified as contamination / host with ContScout and the area under the curve statistics was calculated for each prediction. Matrix of boxplots show distributions of AUC values, grouped according to the source HLTs (corresponding to panel matrix columns) and target HLTs (corresponding to panel matrix rows). Within each of the boxplots, axis x refers to the amount of alien proteins transferred.

### Performance assessment on manually filtered genomes

Contaminant sequences, manually curated from the *Aspergillus zonatus* (filamentous fungus, n=1476)^51^) and *Bombus impatiens* (bumblebee, n=680) proteomes by the authors^15,51,52^, were used as a ground truth when comparing the performance of ContScout with that of BASTA and Conterminator. While ContScout managed to accurately mark all 1,476 manually confirmed bacterial proteins from *A. zonatus*, BASTA identified 1,341 bacterial proteins and Conterminator marked only 948 proteins. None of the three tested tools made any false positive prediction on genuine *A. zonatus* sequences.

For *Bombus impatiens*, overlap between the contaminated insect proteome^52^ and proteins of the independently published symbiont draft genome^15^ were considered as a manually validated collection of contaminating sequences and were used as ground truth. ContScout again found all 680 sequences as contaminants while BASTA marked only 162. Conterminator identified only 8 sequences.

In addition to these examples, we also assessed the performance of the three tools on *Q. suber* proteins containing fungal-specific domains (n=560, see Methods). Out of 560 query sequences, ContScout accurately tagged 556 as fungal contaminants. BASTA detected 44 while Conterminator only identified two of the proteins.

BASTA screening was repeated with a more permissive similarity threshold (50% sequence identity instead of the default value I=80). After relaxing the similarity threshold, Basta accurately called 190 query proteins as fungal contaminants but still missed 390 sequences from the test set (70%). Notably, out of the 44 proteins that were correctly identified as fungal contaminants by Basta with the default threshold, 36 could no longer be identified with the relaxed similarity settings resulting in “Unknown” or “Eukaryote” taxon calls. These results indicate that the classification performance of BASTA highly depends on the similarity threshold set by the user.

### Comprehensive comparison between ContScout and Conterminator

In order to comprehensively compare ContScout and Conterminator, we performed the screening of the entire G844 data set with both ContScout and with Conterminator^13^. BASTA was omitted from this comparison due to the excessive computational resources this tool would require to analyze all 844 proteomes.

Using the Linclust-based protein search mode with default clustering parameters, Conterminator identified only 327 contaminants among 14,749,299 proteins from 844 genomes. In order to provide the tool with the most comprehensive database possible currently, similar to the search conditions applied by Steinegger et al^13^, we repeated the analysis on the non-redundant union of the Uniref100 database with proteins of the 844-genome (See methods for details). In this case, Conterminator marked 18,016 proteins as contaminants including 4,513 hits from the 14.7M proteins from the 844 genomes. At the same time, ContScout marked 51,222 proteins for removal within this set. Comparing the two hit lists we found that 93.24% of Conterminator hits were also reported by ContScout while only 8.22% of ContScout hits were confirmed by Conterminator (Fig. 3). As a coarse proxy for taxonomic affinity, a query-hit HLT match ratio value was calculated based on the top 10 hits of each query protein. More than 99% of the proteins that were marked as contaminants by both software had this ratio below 0.25. Similarly, 98% of proteins that were classified as non-contaminant by both tools displayed at least 75% taxonomy support. It is worth noting that 94% of contaminants that were exclusively predicted by ContScout displayed less than 0.25 taxonomy support, resembling those proteins that were concurrently predicted as contamination by both tools. At the same time, 55% of proteins marked only by Conterminator had at least 0.75 taxonomic support suggesting potential false positive hits (Fig. 3). Taken together, these figures indicate that ContScout outperformed Conterminator, the most up-to-date tool for contamination detection for protein sequences.

**Figure 3:**
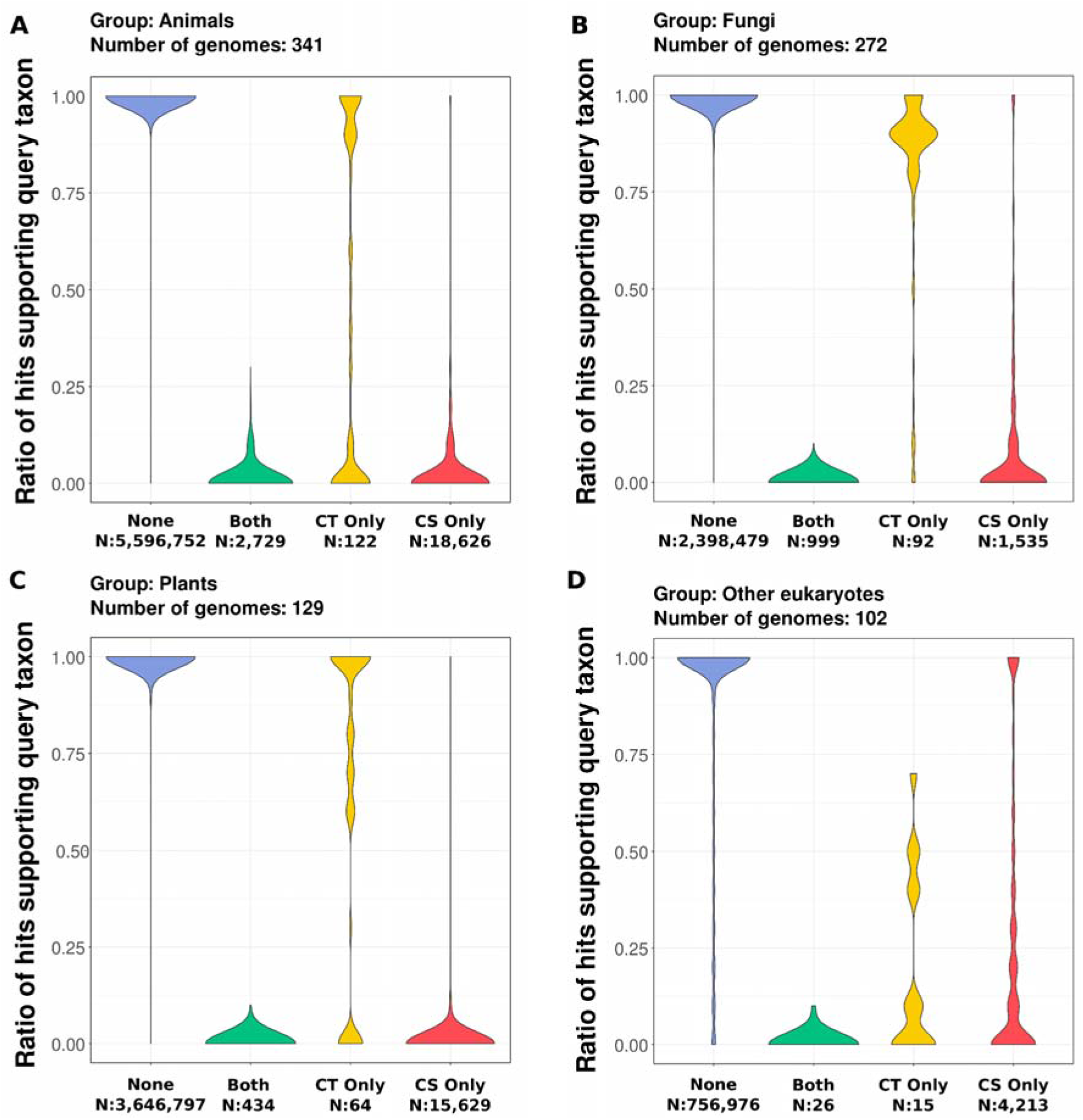
Performance comparison between Conterminator and ContScout. Proteomes of the G844 data were merged non-redundantly with the UniRef100 database enabling a database screen setup similar to what was applied by^13^. ContScout screening was carried out with default parameters as described in the methods section. Proteins were arranged into four groups according to the tools they called them as contaminants: none, both, only Conterminator (CT only) or only ContScout (CS only). For each query sequence a coarse taxonomy support estimator was determined by taking the top 10 hits from the taxonomy-aware UniRef100 database and calculating the rate of hits that confirmed the query HLT. Matrix of violin plots summarize ratios of hits supporting query taxon for each protein group, visualized separately according to the query HLT. (**A**: Animals, **B**: Fungi, **C**: Plants, **D**: Other eukaryotes)

### ContScout detects rampant and diverse contamination in eukaryotic genomes

To assess levels of contamination in public genome databases, we analyzed ContScout outputs obtained on the protein database containing 844 proteomes from all major eukaryotic groups (341 metazoans, 129 plants, 272 fungi and 102 other eukaryotes). ContScout revealed the presence of rampant contamination, detecting at least one contaminating protein in 447 proteomes, reporting 114 contaminants per proteome on average (range: 1-12,656). The presence of contamination in the tested proteomes was smallest among Fungi (43% of the species), slightly higher in Animals (55%) and Plants (56%) and highest among Other eukaryotes (66%).

Bacteria (30,666) and Fungi (17,531) turned out to be the most frequent sources of contamination in the G844 dataset. Of the fungal proteins 12,656 could be linked to a single contaminated plant proteome *Quercus suber* (Fig. 4/B). Viridiplantae (1,538) and Metazoa (1,069) together accounted for no more than five percent of the contaminating proteins. Viruses yielded 273 contaminants while 76 contaminating proteins with Archaea origin were detected.

**Figure 4:**
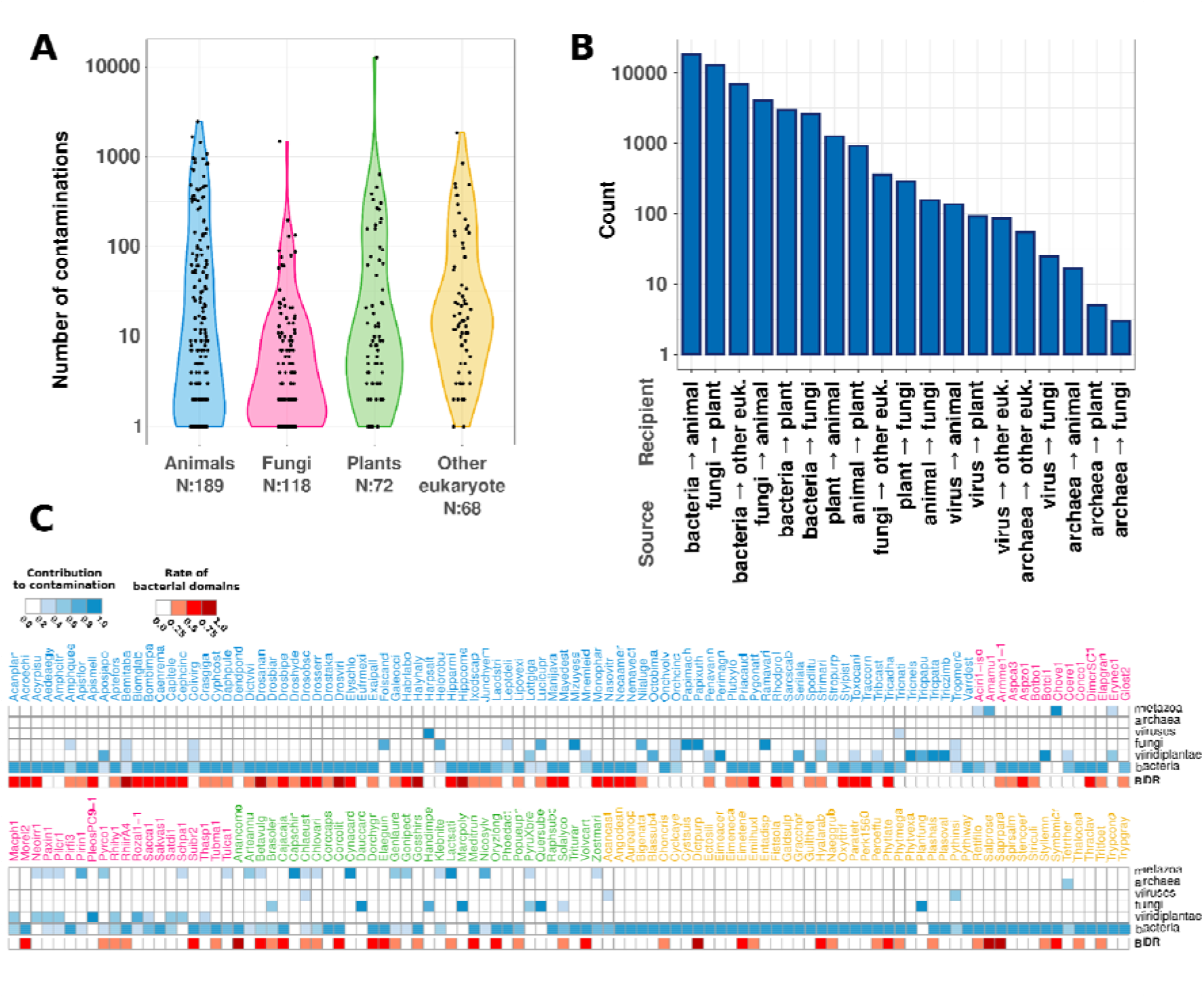
Summary of contamination statistics across 844 genomes. **A**: Violin plot showing the number of contaminant proteins detected in the G844 data set. Proteomes with no contamination (N: 397) were omitted from the plot. Data points were sorted in groups according to the query genome HLT membership. **B**: Barplot summarizing the numbers of contamination-host pairs detected in the G844 data. **C**: Heatmap (blue cells) indicate the contributions of each HLT group to contaminants detected in each of the top 200 contaminated proteomes. Genomes were grouped according to their HLT membership and were ordered alphabetically. Column annotation (red cells) corresponds to the ratio of domains of bacterial origin among the domains detected on contaminant proteins.

From the 200 most contaminated proteomes, 140 had contaminants originating from multiple sources while in 60 cases contaminants could be traced to a single HLT. Bacteria turned out to be the sole source of contamination in 55 proteomes (Fig. 4/C). The prevalence of Bacteria as a source of contamination was also confirmed by Pfam domain analysis within the contaminant sequences: in 52 out of the 200 most contaminated proteomes, Pfam domains exclusive to Bacteria made up more than 50% of the surplus domains that were called on the contaminant sequences. Sequences marked by ContScout are listed in Supplementary table 1.

### Phylogenomic analyses are biased by contamination

We next addressed the hypothesis that contamination might bias phylogenomic analyses of gene content and reconstructions of ancestral genomes. Ancestral genome reconstructions have recently gained momentum in several organismal groups and have revealed broad patterns of genome evolution. For example, an early burst of gene duplication followed by gene loss appeared as a dominant mechanism of genome evolution in the metazoa, land plants and fungi (e.g.^31,33,53–57^. However, how contamination influences these analyses has not been assessed to date. We addressed this question here using reconstructions of ancestral gene content in early eukaryote ancestors, including the last eukaryotic common ancestor (LECA), using original and decontaminated data.

We used a 36-species data set containing 10 contaminated genomes and inferred gene gain, duplication and loss patterns using the original genomes and those cleaned by ContScout. Figure 5 shows considerable differences in inferred ancestral genome sizes between contaminated and clean data. For example, based on contaminated data, LECA possessed 10,764 protein-coding genes, whereas in decontaminated data it possessed 8,919, a 20.6% overestimation. The largest difference (88% overestimation) was observed in the Opisthokonta MRCA (N49 on Figure 5), in which contaminated data suggested 6,097 more ancestral genes (14,216) than decontaminated data did (8,119). This node reflects a combined signal from multiple contaminated genomes, including fungal genes in the cork oak (12,650 genes) and bacterial genes in the bumblebee (965 genes). This is clearly an extreme case due to the inclusion of two massively contaminated genomes, however, it reveals that contamination behaves additively as we move from recent to ancient nodes in the tree. It is thus conceivable that significant overestimation of ancestral gene counts can build up from combined effects of even moderately contaminated genomes. It is also noteworthy that contamination affected ancestral gene numbers along the entire phylogenetic path that connects contaminants with the contaminated genome.

**Figure 5:**
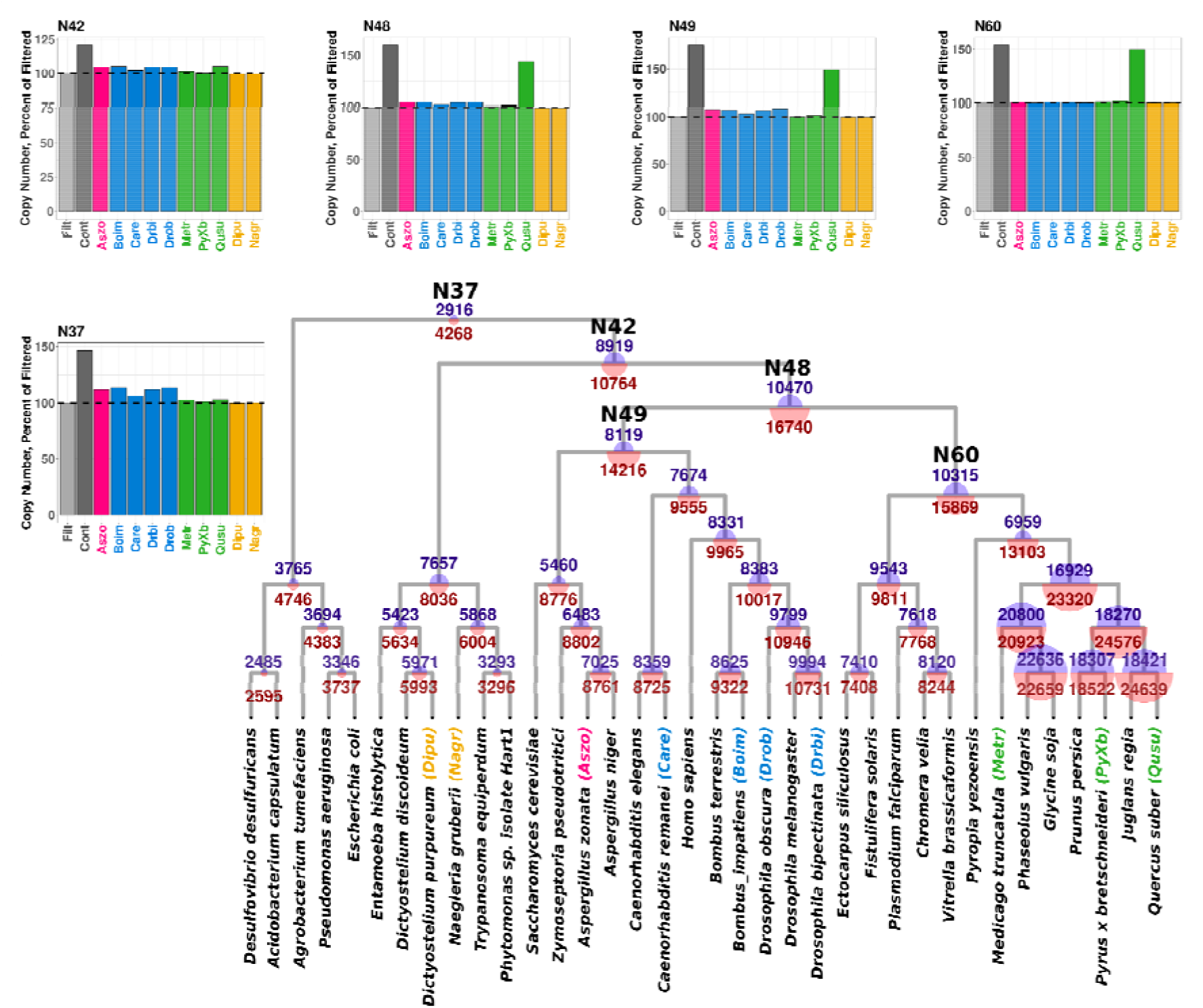
Effect of contamination on ancestral genome reconstruction in the 36G data set. Copy number estimates are provided as internal node labels within the species tree. Estimates based on contaminated data are printed in red while values inferred from clean data are colored blue. Bar plots show bias of individual contaminated genomes introduced to copy number prediction at prominent nodes. Barplots show copy numbers as percentages where 100% corresponds to the copy number inferred from clean data at any given internal node. Abbreviations used on barplot legends: **Filt**: decontaminated, **Cont**: contaminated, **Aszo**: *Aspergillus zonatus*, **Boim**: *Bombus impatiens*, **Care**: *Caenorhabditis remanei*, **Drbi**: *Drosophila bipectinata*, **Drob**: *Drosophila obscura*, **Metr**: *Medicago truncatula* **PyXb**: *Pyrus x bretschneideri*, **Qusu**: *Quercus suber*, **Dipu**: *Dictyostelium purpureum*, **Nagr**: *Naegleria gruberii*. Prominent internal nodes: **N37**: LUCA, **N42**: LECA, **N48**: Opisthokont + Archaeplastida MRCA, **N49**: Opisthokont MRCA, **N60** TSAR + Archaeplastida MRCA

Ancestral genes inferred in a node due to contamination displayed a clear functional signal for the source. The 1,845 genes that were only inferred in LECA in the contaminated data set were clearly enriched in protein domains occurring mostly or exclusively in bacteria (Supplementary Table 2). Such alien functions in ancestral genomes can clearly distort our functional understanding of ancestral species. It is worth noting that we detect bacterial contamination (n=60) in the *Naegleria* genome, which was used to infer an ancestral gene content of eukaryotes^26^, and could potentially also contribute to an ancestral gene content overestimation problem.

Across the entire data set, the contaminated analysis suggested 11,029 more gene gains (*de novo* origins and duplications) and 56,687 more gene losses than the decontaminated analyses, indicating that both metrics are biased by contamination, but gene losses are affected nearly five times more. Using a series of selectively decontaminated data, we separately measured the effect of contaminants from each of the ten contaminated genomes within the G36 data set. As expected, ancient gene copy estimation overestimation at the MRCA of Archaeplastida / TSAR (N60) can be solely explained by contaminations in *Quercus suber* with considerably smaller contribution from other genomes. On the other hand, overestimations at LUCA (N37) mainly originate from insect and fungal genomes contaminated with bacterial sequences. It is worth noting that the effect of individual contaminant genomes clearly sum up and jointly bias the copy number overestimations at LECA (N42) as well as at the opisthokont MRCA (N49).

### Contamination severely inflates gene loss rates

To uncover why contamination inflates gene loss nearly five times more than gene gain estimates, we manually checked gene families that include contamination. For illustrating the mechanism, we chose the pyridoxal kinase protein family that is part of the pyridoxal 5’-phosphate salvage pathway, is conserved across the entire tree of life and ContScout identified one contaminating proteins in *Q. suber* and one in *B. impatiens* (Supplementary Fig. 2). The maximum likelihood gene tree readily identified two mis-positioned proteins. One of the *Q. suber* proteins “(Quersube_4764)” clustered with fungal pyridoxal kinases (SH support value: 0.98), whereas one from *B. impatiens* (“Bombimpa_11962”) was clearly positioned in the bacterial clade (SH support value: 0.97). Both suspicious proteins were tagged as contamination by ContScout together with further 94 and 60 other proteins in *Q. suber* and *B. impatiens*, respectively, that were located on the same assembled scaffolds. In line with the gene tree, “Quersube_4764” was tagged as fungal by ContScout while the tool predicted “Bombimpa_11962” to originate from bacteria.

We performed reconciliation and mapping of gene gain/loss events for this protein family on the species phylogeny, using the method we used for the genome-wide mapping. This indicated that the contaminated and decontaminated gene trees can be explained by 26 (6 gene gains and 20 losses) and 8 (7 gains, 1 loss) events, respectively (Fig. 6).

**Figure 6:**
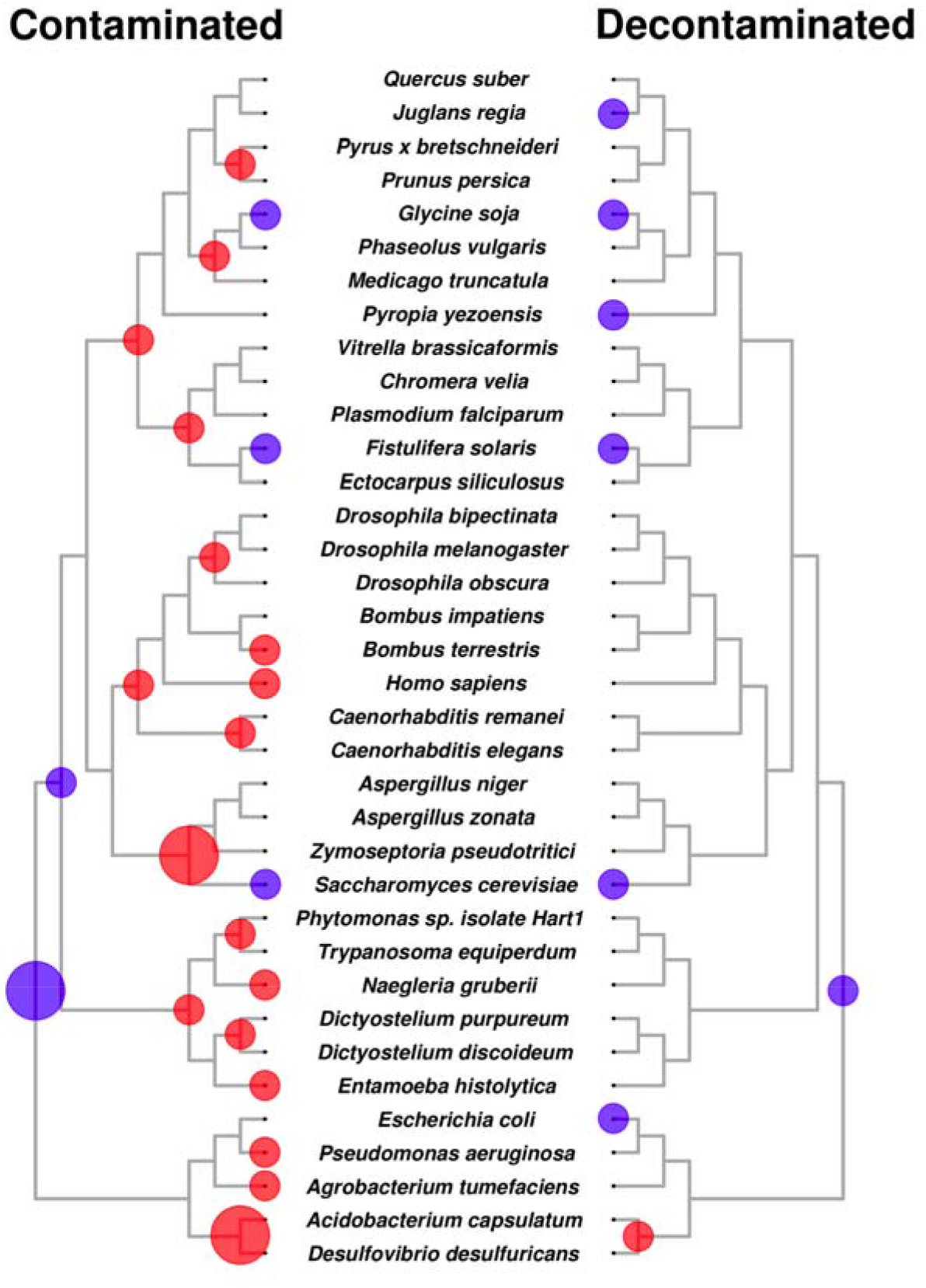
Bias of contamination introduced to gene loss / gene gain inference demonstrated by the case study of the pyridoxal kinase protein family. Gene family evolutionary events that were inferred based on contaminated data are mapped in the left part of the figure while the right part contains inferences based on clean data. Blue circles indicate gene gains while red circles correspond to gene losses. At each node of the tree, circle sizes are proportional to the number of mapped events.

LECA had one versus two ancestral genes in this family in the decontaminated and contaminated analyses, respectively. This trend is similar to the genome-wide numbers (see above) and is a significant difference considering that the contaminated and decontaminated gene trees differed only in two proteins. If those two proteins were both species-specific paralogs, as, for example 19577 and 27563 of *Juglans regia* (walnut), then they would have necessitated only two more events. However, we found that Quersube_4764 and Bombimpa_11962 have induced eight and ten gene losses, respectively. In the case of Quersube_4764 these losses were introduced because, during the mapping, Quersube_4764 was assigned to a 1-to-1 orthogroup with a fungal protein (Zymps_805618) and due to the definition of orthology, the origin of the orthogroup [Quersube_4764, Zymps_805618] had to be in the most recent common ancestor of plants and fungi. It follows that for this orthogroup to be explained along the phylogeny, losses had to be counted for all descendents of the plant/fungal ancestor except *Q. suber* and *Z. tritici*.

We define here orthogroups as sets of genes that are separated from each other by speciation events, as used by most reconciliation-based gene family analyses (^22,58–60^. Several studies define orthogroups more loosely, as sets of similar proteins that cluster together in a similarity-based clustering approach (e.g. MCL (^61,62^). We expect these simpler approaches to be less sensitive to distorting effects of contamination, simply because they don’t resolve gene family evolution at the level of individual genes’ duplication and loss events. Yet, such studies can still be heavily affected by contamination. At the level of the 36-species data set, we identified 1,827 and 259 clusters that consist entirely of fungal + *Q. suber* or bacterial + *B. impatiens* proteins, respectively. The *Q. suber* and *B. impatiens* proteins of 1,814 and 259 of these clusters have been identified by ContScout as contamination. In the phylogenetic mapping, these clusters contributed 13,043 and 2,223 losses, respectively, and pushed gene family origins earlier, even if orthogroups are equated with MCL-inferred similarity-based clusters.

## Discussion

In this paper we presented a new method for identifying contaminating proteins in assembled genome sequences and demonstrated that in evolutionary genomics contamination can lead to a false notion of complex ancestral genomes and high gene loss rates. Contamination is a widely recognized problem in sequence databases and can stem from a variety of reasons (reviewed by Cornet et al.^63^). Several tools have been developed for the detection of contamination in large sequence databases^13^ or estimating contamination level in (meta)genomes (e.g. CheckM^36^, Busco^37^). Most previous tools focus on classifying raw sequencing reads as host or alien^39,42^ or rely on measures of similarity to a pre-selected set of marker genes or ribosomal proteins^36,37,64^, whereas genome-wide tools that can clean genomes of contamination are at paucity^13^. ContScout is a genome-wide method that relies on a reference database and genome annotation data to identify and remove contaminating proteins. After inferring protein-wise taxon calls similarity searches against taxon-aware reference databases, ContScout summarizes these across contigs/scaffolds and flags sequences which appear to be contamination. Thus, ContScout can clean genomes of contaminating proteins and yield clean genomes. We anticipate this feature will become more and more important as genome sequencing efforts are extended to field and museum specimens, mixtures of organisms (e.g. host and its parasite, metagenomes) or unculturable single-cells, all of which increase the risk of contamination. Our analyses of synthetic data, benchmarking against manually curated sequences, as well as a comparison to other decontamination tools of protein sequences indicated that ContScout achieved high sensitivity and specificity. We think this is rooted in the combination of RLE-based taxon calls with scaffold-level decision making. First, the automatic RLE method may yield more robust taxon call assignment than any fixed-size hit lists (e.g. top 100 hits) and can help minimize the effect of mis-labeled proteins, sporadically present in reference databases. Second, if a contig is marked for removal by ContScout, any ambiguous proteins coded on it will be also discarded, which potentially increases sensitivity. On the other hand, horizontally acquired genes, which are integrated into host chromosomes, are conceivably not discarded by ContScout, because the majority of proteins encoded on those chromosomes/scaffolds will be classified as host.

We found that ContScout greatly outperformed Conterminator and BASTA, two recent cleaning tools operating in the protein space. For example, while ContScout identified all proteins that were manually flagged as contamination in *A. zonatus* ^51^, Conterminator and BASTA identified only 64% and 91% of them, respectively. We hypothesize that the loss sensitivity of both BASTA and Conterminator lies in their search logic (i.e. using fixed similarity thresholds with values being set rigorously high by default) that implicitly assumes that the contaminating organism (or its close relative) is present in the reference database. Even with the dense sampling of genomes we have today, having the genome of the exact contaminating taxon in the database is a rare situation, so we think the more dynamic and sensitive search engine implemented in ContScout is warranted.

ContScout’s limitations, on the other hand, lie in decisions on draft genomes with low N50 values (i.e. fragmented assemblies with small contigs), screening within undersampled groups (e.g. protists) and in chimeric contigs, the latter of which albeit exist, are likely rare^65,66^. The current implementation of ContScout focuses on protein-level detection of contamination across large taxa, such as fungi, plant and prokaryotes, however, its search logic can be easily extended to finer taxonomic scoring, to DNA sequences and to search among prokaryotic taxa. While the latter is realistic given the broad availability of prokaryotic reference genomes, assessing contamination by a taxon to another in the same eukaryotic kingdom may currently be limited by the paucity of genomes in several eukaryotic groups. Using a finer taxonomic scoring would be justified for improving ContScout’s ability to detect human contamination from metazoan genomes, which is currently limited.

Using ContScout we screened 844 published eukaryotic genomes and found widespread contamination by various organisms, most commonly bacteria and fungi. We identified >50,000 contaminating proteins in this set, while Conterminator identified 327 - 14,148 depending on the database configuration used (screening within G844 alone or G844 united with UniRef100). These figures agree with previous reports of contamination in reference sequence databases^67^ and (meta)genomes^12,36,68–70^, however, our inventory highlighted a range of novel patterns. First, the number of contaminating proteins covered three orders of magnitudes: it ranged from a handful of proteins up >14,000, in extreme cases allowing the subtraction of presumably complete protein repertoires of the contaminating organism^14,15^ from the contaminated genome. Concomitantly, the sizes of scaffolds encoding contaminant proteins reached 1 Mb (*A. zonatus*). This diversity of contamination depths challenges tools that use fixed parameters (such as contig lengths^13^, similarity thresholds^13,48^ or hit lists^41^, justifying the relative cutoff-free approach ContScout uses. Second, the taxonomic distribution of contaminating organisms reflects common lifestyles of microbes as symbionts, parasites, food sources or commensals. Previous studies have also reported bacteria as a common contaminant^8^ whereas in our analyses fungi also emerged as frequent contaminating organisms, possibly due to their diverse associations with plants and metazoans. Finally, we expect the cleaned genome annotations for 844 eukaryotes, covering all eukaryotic supergroups and phyla can form a gold standard resource on which comparative analyses can be built.

Whereas the effects of contamination on metagenomic studies and functional interpretation of genomes is quite straightforward^41,65^, the biases they cause in the context of evolutionary genomics is poorly explored. Such studies hold the promise of looking back in time and identifying the genomic blueprint of major radiations. We found that such studies can suffer considerably when contamination in the data set: alien proteins pushed gene family origins towards the root of the tree, resulting in an overestimation of ancestral gene contents, and induced large numbers of gene losses. This effect was additive when multiple contaminated species were present in the data set, resulting in increasingly more biased estimates towards ancient nodes. This can give the impression of highly complex ancestral genomes, as inferred in recent empirical studies on ancestral gene content in several groups, such as animals, plants^53^ or LECA^26^. A recent report demonstrated that incomplete genome annotation can also inflate gene loss estimates^71^, however, while in that case excess losses are distributed randomly across the tree and affected mostly terminal branches, excess loss introduced by contamination yielded profoundly different distribution of bias, mostly affecting deep branches and causing a several-fold overestimation.

We used early eukaryotic nodes as a case study to assess the impact of contamination on ancestral gene content estimation. Previous studies conjectured that LECA was a complex organism, with at least 4,000^26^, 10,000^72^ or 12,000^73^ genes and that it already possessed several key eukaryotic traits, such as complex cytoskeletal systems and sexual reproduction^26,74–77^

Our study suggests that LECA possessed ~8,700 genes, however, we also show that genome choice can severely influence the inferred complexity of LECA’s genome and may lead to the overestimation of its functional repertoire and the inference of alien functionalities. Broadly, these figures are consistent with a complex LECA genome, however, also draw attention to considering contamination as a source of bias in ancestral gene content estimation.

In summary, our data suggest that in phylogenomic studies false inference of complex ancestral genomes, incorrectly inferred early origins of gene families and an overestimation of losses might go hand in hand with contamination. These, combined with or other sources of bias, such as annotation errors^71^ and unrecognized HGT^76^ can lead to false inferences of ancestral gene content, influencing our perception of the evolutionary process. We expect that ContScout and the analyses presented here will facilitate the accumulation of high quality genomes and improve evolutionary genomics studies for reconstructing how life evolved on Earth based on genomic data.

## Supporting information

List of Supplemental Files

## Acknowledgements

This work was funded by the Momentum Program of the Hungarian Academy of Sciences (LP2019-13/2019) and by the European Research Council (Grant No. 758161) (both to LGN). This research was performed under the Facilities Integrating Collaborations for User Science (FICUS) program (proposal: 10.46936/10.25585/60008430) and used resources at the DOE Joint Genome Institute (JGI) (https://ror.org/04xm1d337) and the National Energy Research Scientific Computing Center (NERSC) (https://ror.org/05v3mvq14), which are DOE Office of Science User Facilities operated under Contract No. DE-AC02-05CH11231.

## Methods

### Selection of the G844 data set

In total, 844 published genomes, comprised of 341 animals, 272 fungi, 129 plants, and 102 other eukaryotes, were downloaded from public databases (JGI MycoCosm^5^, ENSEMBL^78^, Genbank^79^) to perform a broad contamination screening. If multiple isoforms of the same gene were present, we selected the longest one for analysis. Genomes included in the study are summarized in Supplementary table 3. Date of data collection: July, 2019.

### Selection of the G36 data set

In order to assess the effect of contamination on ancestral genome reconstruction, a 36-genome data set has been compiled encompassing five bacteria, eight animals, four fungi, seven plants and twelve other eukaryotes. Altogether, the data set included ten genomes (*Aspergillus zonatus*, *Bombus_impatiens*, *Caenorhabditis remanei*, *Dictyostelium purpureum*, *Drosophila bipectinata*, *Drosophila obscura*, *Medicago truncatula*, *Naegleria gruberii*, *Pyrus x bretschneideri*, *Quercus suber*) that contained between 28 and 12,656 contaminant proteins each. Contaminated genomes, each matched with a contamination-free related genome, were selected so as to represent all major eukaryotic lineages in which we frequently found contamination.

### Domain analyses

For the G844 data set, Interproscan v5.44.79.0^80^ was used to search for protein domains. Bacterial-specific domains were extracted from the Pfam database (Mistry et al., 2021; v 35.0) based on the ratio of bacterial sequences within the seed alignments. Domains with over 95% bacterial sequences in their seed alignments were considered as bacterial. In order to collect Fungi-specific domains, the UniprotKB database together with IPR annotations was downloaded. A domain was considered as fungi-specific if at least 95% of the associated UniProtKB proteins originated from the kingdom Fungi.

### ContScout run parameters

ContScout runs were carried out using the docker image h836472/contscout_avx2. Uniref100 database (release 2022_1) was selected as the reference database (-d uniref100) with MMSeqs used as the search engine (-a mmseqs) with the search sensitivity set to “very fast” (-s 2). The minimum sequence identity threshold was set to 20 percent (-p 20). Contamination from all possible high level taxa (i.e. archaea, bacteria, plant, fungi, animal, other eukaryote, referred in the rest of the article as HLT) was screened (-x all).

### Performance assessment of ContScout on synthetic data

Seventeen genomes (5 bacteria, 4 animals, 3 fungi, 3 plants, 2 other eukaryotes) from the G36 data set with no evidence of contamination were collected and used to assess the contamination / host classification performance of ContScout. Artificially contaminated genomes were created by transferring proteins between pairs of “source” and “recipient” genomes, covering all possible non-identical source / recipient genome HLT combinations.

For each genome pair, a set of 100, 200, 400, 800, 1,600 or 3,200 randomly selected proteins were transferred, assigned to random virtual contigs each holding one, two, five, ten or twenty alien proteins. At each of the six spike-in levels, 100 random replicate sets were generated. ContScout was then executed on the artificially contaminated data to classify proteins as either host or contamination. Area under the curve statistics, calculated by the pROC^81^ package in R^82^, were used to assess classification performance.

### Performance assessment of ContScout on manually curated data

The 844-genome data set included two projects (*A. zonatus*^51^ and *B. impatiens^52^*) where the authors carried out manual genome decontamination and released clean assembly versions after our data collection took place. In addition, Martinson and co-workers released a separate draft genome for the gut symbiont gamma proteobacterium^15^, which they identified as the sole source of contamination within the original *B. impatiens* assembly. For *A. zonatus*, we used the proteins that were removed by authors as a ground truth. For *B. impatiens* we mapped back proteins from the published symbiont proteome to the contaminated *B. impatiens* data using a sequence identity threshold of 95% and a sequence coverage threshold of 0.6. That way, we identified 680 symbiont proteins within the *B. impatiens* proteome that served as positive control for contamination. We used these protein sets to compare the performance of ContScout to that of Conterminator and BASTA.

The *Q. suber* proteome was manually screened for the presence of fungal contamination using Interpro domains that were found specific to fungi within the UniprotKB database. That way, we identified 560 protein sequences that are very unlikely to be part of the *Q. suber* proteome. We then compared the sensitivity of ContScout, Conterminator and BASTA using these strong *Q. suber* contaminant candidates.

### Large-scale comparison between ContScout and Conterminator

In order to carry out a Conterminator screen similar to that of^13^, the 844G data set was combined with the UniRef100 database (release 2022_01). Whenever a protein sequence was present in both sources, redundancy was resolved by keeping only the copy from the 844-genome set. Conterminator was then executed in “protein” mode, using default parameters. Proteins of the 844-genome set were sorted into four classes according to hits lists of the two compared tools: 1., tagged by neither software, 2., tagged by both tools, 3., tagged only by Conterminator, 4., tagged only by ContScout.

Independently, each query sequence was aligned against the UniRef100 database keeping the ten best-scoring hits per query. Ratios of hits from the same high level taxon (HLT) as the query protein were calculated as an indicator of taxonomic support and were visualized separately for each results class. A total of 146 proteins that were marked exclusively as contamination either by ContTerminator (4) or ContScout (142) orthogroup information together with a species tree was available in the G36 data set; therefore we manually inspected the position of those proteins within the gene family phylogenies, as validation.

### Ancient genome reconstruction and gene copy number estimation

We followed published pipelines for reconstructing ancestral genomes using the G36 dataset which, briefly, utilized a species tree and reconciled gene trees for each of the protein families identified in the input genomes^22,53,55^. For inferring a species tree, BUSCO^37^ v3 HMM profiles were used to collect 428 conserved single-copy candidate proteins from the decontaminated G36 data set. MMSeqs was then applied to calculate an all versus all protein similarity network among the proteins on which Markov-clustering with hipMCL^83^ was carried out with an inflation parameter of I=2 to identify protein families. Predicted protein families were filtered manually, keeping only the conserved single-copy ones. Mafft (v7.407^84^) with the “--auto” option was used to perform multiple sequence alignment for each single-copy protein family. Uninformative and poorly aligned parts were removed with TrimAl^85^ (parameters: “-gt 0.95”) and the resulting trimmed alignment were concatenated into a supermatrix of 428 protein families and 172,083 characters. RAxML 8.2.12^86^ was used to infer a maximum-likelihood species tree under the PROTGAMMALG model of protein evolution. The model was partitioned gene-by-gene.

For assessing the impact of contamination on ancestral genome reconstruction, a series of semi-decontaminated data sets was generated based on the G36 collection where each dataset retained contamination from only one of the ten contaminated genomes (*Aspergillus zonatus*, *Bombus_impatiens*, *Caenorhabditis remanei*, *Dictyostelium purpureum*, *Drosophila bipectinata*, *Drosophila obscura*, *Medicago truncatula*, *Naegleria gruberii*, *Pyrus x bretschneideri*, *Quercus suber*). The data series was then completed with the fully decontaminated as well as the original G36 versions. For each variant in the series, orthologous protein families were identified by Orthofinder v2.4.1^58^ using the species tree as a reference. Ancient genome reconstruction as well as gene gain / loss events were inferred by using the COMPARE pipeline as described before^22^. Effects of individual contaminated genomes were determined by comparing gene gain / loss counts between contaminated and clean versions.

## Software availability

ContScout tool can be downloaded as a Docker image from repository *h836472/contscout_avx2*. Source code for ContScout is available at *https://github.com/h836472/ContScout/*

## Notes

### Competing Interest Statement

The authors have declared no competing interest.

