## Supplementary material for "Purging genomes of contamination eliminates systematic bias from evolutionary analyses of ancestral genomes": List of Supplemental Files

**List of contents**

**Supplementary table 1: (Supplementary_table_1_CS-Contams.xlsx)** Sequences marked by ContScout as contamination in the G844 data set.

**Supplementary table 2: (Supplementary_table_2_LECA_IPR.xlsx)** IPR domains significantly enrichment in proteins that were excessively predicted as part of the ancient genome of LECA as a result of contamination in the G36 data set.

**Supplementary table 3: (Supplementary_table_3_G844_database.xlsx)** Details of the 844 eukaryote genomes used in the study.

**Supplementary figures:** (**SupplementaryFigures.docx)**
