## Supplemental Figures for "Purging genomes of contamination eliminates systematic bias from evolutionary analyses of ancestral genomes"

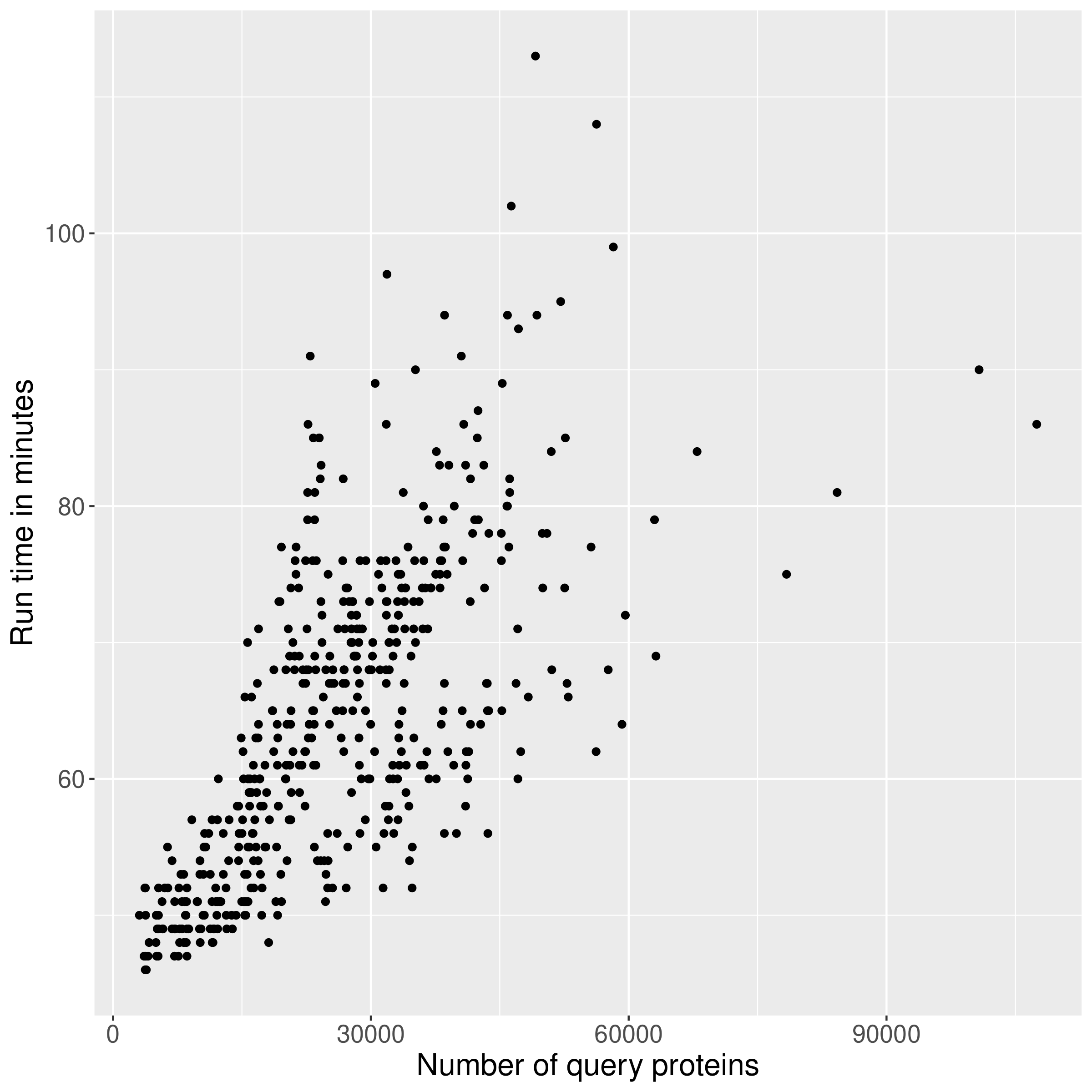

**Supplementary Figure 1**: ContScout run times measured plotted as a function of query proteome size. Runs were performed using 24 CPU cores with RAM usage constrained to 150 GB.

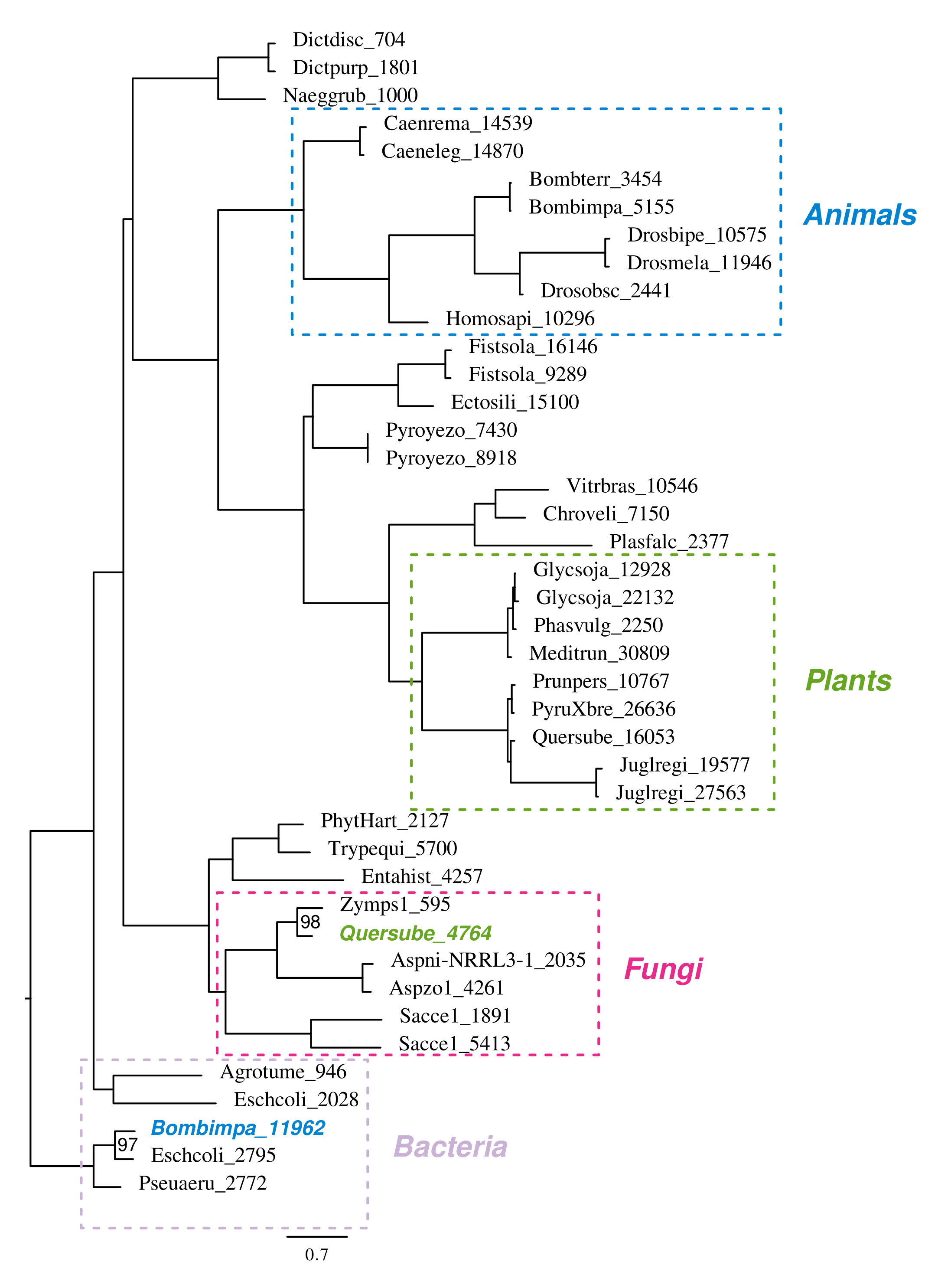
**Supplementary Figure 2**: Gene tree of the pyridoxal kinase protein family based on the unfiltered G36 data. Dotted lines indicate high level taxons (Animals in blue, Plants in green, Fungi in magenta and Bacteria in pale purple.) The Positions of *Q. suber* protein *Quersube_4764* and *B. impatiens* protein *Bombimpa_11962* clearly indicate fungal and bacterial contamination, respectively.
